## Supplementary Text, Figures, and Tables for "Overcoming bias in gene-set enrichment analyses of brain-wide transcriptomic data"

#### S1 Results

##### S1.1 GO Category Case Study

To demonstrate the mechanisms underlying the major differences between randomized-gene nulls and randomized-phenotype nulls, especially in the case of spatially embedded (and spatially autocorrelated) transcriptional atlas data, we investigated the null distributions of three specific GO categories in detail. We selected two similarly-sized GO categories in mouse that have different levels of within-category coexpression: 'regulation of dendritic spine morphogenesis' (RDSM, 40 genes), and 'zymogen activation' (ZA, 42 genes).

###### S1.1.1 Within-category coexpression drives false-positive bias

We first aimed to understand why, under the SBP-random ensemble, RDSM (CFPR = 13%) has a higher CFPR than ZA (0.07%). Region × gene expression matrices and gene × gene coexpression matrices are plotted for RDSM and ZA in Figs S3A and B, respectively. Compared to a representative random set of 40 genes, shown in Fig. S3C, we find a much more consistent spatial patterning of gene expression in the RDSM category, resulting in increased within-category (gene–gene) coexpression. This spatial coherency of expression patterning is consistent with genes associated with regulating dendritic-spine morphogenesis (RDSM) having a coordinated brain function that varies characteristically across brain regions. By contrast, GO categories that play a minimal or less-specific role in brain function, such as zymogen activation (ZA), exhibit noisier expression patterns and lower within-category coexpression (that is visually similar to that of a random set of genes).

To understand how these differences in within-category coexpression affect category false-positive rates, we investigated the distribution of mean correlation coefficients between the expression profiles of genes in each category and the ensembles of randomized SBPs analyzed above. These category-score distributions are plotted for SBP-random and SBP-spatial ensembles in Figs S3D and E, respectively. A category with wider distributions than a random set of genes (the null comparison used in GSEA) will have a greater probability of obtaining a significant correlation to that ensemble of phenotypes. First we note that, consistent with both SBP-random and SBP-spatial ensembles containing no information about gene expression, all category-score distributions are symmetric about zero.

Under random phenotypes (Fig. S3D), RDSM has a wider distribution of category scores than ZA; it is more likely to exhibit a higher correlation to a random SBP. This widening is driven by the increased coexpression of RDMS genes, such that a chance correlation between a random SBP and any single RDMS gene is amplified through a similar correlation with many other genes in the category. By contrast, categories like ZA contain genes with lower gene–gene coexpression, such that a chance correlation between an SBP and a given ZA gene is more likely to be cancelled out by a chance correlation in the opposite direction for another gene, driving category scores towards zero and resulting in a narrower category-score distribution.

In estimating category $p$-values, conventional GSEA compares the category score to that of a random set of the same number of genes, shown gray in Fig. S3D (where we have performed many randomizations of sets of 40 genes so as not to bias towards any particular random set). Due to the high coexpression of RDSM genes, this occurs for approximately 13% of SBP-random phenotypes, but for ZA, which has a similar coexpression structure as random genes, this occurs for just 0.07% of random SBPs.

###### S1.1.2 The role of spatial autocorrelation

Relative to the SBP-random ensemble (of purely random phenotypes), the CFPR of RDSM increased under the SBP-spatial ensemble: 13% $\rightarrow$ 28%, whereas the CFPR for ZA decreased: 0.07% $\rightarrow$ 0.02%. Examining the category-score distributions in Fig. S3E, we first note that they are all much wider than for the SBP-random ensemble (Fig. S3D), including for random sets of genes. This is due to the predominance of spatial autocorrelation in the expression patterns of individual genes: a gene that exhibits a strongly autocorrelated expression map is more likely to be strongly correlated to a spatially autocorrelated SBP than a random spatial map. Relative to random genes, we see a wider category-score distribution for RDSM, and a narrower distribution for ZA, consistent with their CFPRs. Imposing the constraint of spatial autocorrelation (i.e., SBP-rand $\rightarrow$ SBP-spatial) can thus either increase a GO category's CFPR (if it exhibits a more similar spatial autocorrelation structure to the SBP-spatial ensemble) or decrease it (if it exhibits a less similar spatial autocorrelation structure).

#### S2 Phenotype Enrichment

Full GO enrichment results for all phenotypes across all three null models are in supplementary files. For human cortex:

- Node structural connectivity betweenness, $B$, `EnrichmentThreeWays_human_cortex_betweenness.csv`
- Node structural connectivity degree, $k$, `EnrichmentThreeWays_human_cortex_degree.csv`

And for mouse (whole-brain and cortex), where all files share the `EnrichmentThreeWays_mouse` prefix; suffixes are given below:

- Node structural connectivity betweenness: `_all_betweenness.csv`, `_cortex_betweenness.csv`
- Node structural connectivity degree: `_all_degree.csv`, `_cortex_degree.csv`
- VIP+ cell density: `_all_VIP.csv`, `_cortex_VIP.csv`
- SST+ cell density: `_all_SST.csv`, `_cortex_SST.csv`
- PV+ cell density: `_all_PV.csv`, `_cortex_PV.csv`
- oligodendrocyte density: `_all_oligodendrocytes.csv`, `_cortex_oligodendrocytes.csv`
- neural density: `_all_neurons.csv`, `_cortex_neurons.csv`
- glia density: `_all_glia.csv`, `_cortex_glia.csv`
- microglia density: `_all_microglia.csv`, `_cortex_microglia.csv`
- excitatory density: `_all_excitatory.csv`, `_cortex_excitatory.csv`
- inhibitory density: `_all_inhibitory.csv`, `_cortex_inhibitory.csv`

#### S3 Literature Survey

Enrichment on the basis of spatial patterns of expression has not been performed consistently in the existing literature. Studies have processed the data differently (including substantially different methods for normalizing and filtering genes), and performed the enrichment differently (using different software packages, annotation systems, and including different sets of categories for enrichment). Our survey focuses on studies that have reported gene-set enrichment for Biological Processes in the Gene Ontology (GO) [3].

#### S3.1 Allen Mouse Brain Atlas

The following studies report results of GSEA using transcriptional data from the Allen Mouse Brain Atlas (AMBA) [4]:

1. French et al. [5] (mouse expression and rat connectivity) 3976 coronal genes using ORA (*ermineJ* [6]) for the two anti-correlated expression patterns: NE (neuron-enriched pattern) and OE (oligodendrocyte-enriched pattern). Data taken from Supplementary Data Sheet 3.

2. French and Pavlidis [7] (mouse expression and rat connectivity) 17 530 genes using ORA (*ermineJ* [6]). Data taken from proximity-corrected enrichment for (i) outgoing connectivity, and (ii) incoming connectivity (Table S5). [[check that actual analysis derives from here]]

3. Ji et al. [8]: Tabulated enrichment categories that appeared often across many analyses using 4084 coronal section genes: all brain structures and different ways of measuring connectivity. Data taken from Table 1.

4. Fakhry and Ji [9]: 4084 (coronal-section) genes that are predictive of voxel-level brain connectivity using ORA. Data taken from Fig. 5, summarizing each GO category as the number of injection structures for which it was significant.

5. Rubinov et al. [10]: 3380 genes (assayed multiple times) scored by partial least squares for nodal participation metrics, doing enrichment on the genes in the top 25% of positive weights. Data taken from Table S3.

6. Fulcher and Fornito [11]: 17 642 genes scored by differences in gene coexpression contribution (GCC) scores using GSR using *ermineJ* [6]. Two GSEA were performed, for: (i) connected versus unconnected pairs of brain areas (data taken from Table S1), (ii) rich and feeder connections versus peripheral connections (data taken from Table S5).

7. Mills et al. [12]: 3079 (coronal-section) genes, enrichment for processes showing a strong relationship between CGE and functional connectivity (FC) using ORA (*ermineJ* [6]). Data taken from Table 3.

8. Ko et al. [13]: 170 neuron-, 44 oligodendrocyte-, and 50 astrocyte-specific genes for the coronal plane (gene sets taken from [14] using a 10-fold threshold, and including only genes that could be matched to AMBA data). Enrichment performed on each gene set, with results summarized in the text (Page 2).

#### S3.2 Allen Human Brain Atlas

The following studies report the results of GSEA using data from the Allen Human Brain Atlas (AHBA) [15]:

1. Tan et al. [16]: 20 444 genes, ORA enrichment (using *DAVID* [17]) on top 100 genes for the most positive and negatively correlated expression patterns (one marks neurons, the other for oligodendrocytes). Data taken from Table S6 ($p$-values for 'positive 100') and Page 8 text (list of significant categories for 'negative 100').

2. Richiardi et al. [18]: 16 906 genes: enrichment on strength of correlated gene expression within functional networks (relative to between functional networks) performed with *DAVID* [17] and Panther [19]. Data taken from Table S6.

3. French and Paus [20]: 20 737 genes, ROC enrichment (ermineJ [6]) for measures of inter-subject consistency and inconsistency. Data taken from supplementary files from `https://figshare.com/articles/A_FreeSurfer_view_of_the_cortical_transcriptome_generated_from_the_Allen_Human_Brain_Atlas/1439749`: `InconsistentGOGroups.tsv` and `ConsistentGOGroups.tsv`.

4. Vértes et al. [21]: 20 737 genes, enrichment on topologically integrative hubs (of functional connectivity), for positive and negative contributions to partial least squares components 1, 2, and 3. Data from supplementary file: `Vertes-rstb20150362supp1.xlsx`.

5. Parkes et al. [22]: 19 343 genes, enrichment performed using GSR (ermineJ [6]) on coefficients of principal components (PCs) of gene expression in the striatum, for PCs 1, 2, 5, and 9. Data obtained directly from the author.

6. Forest et al. [23]: 20 783 genes, enrichment performed on gene clusters formed using WGCNA [24] using *Cytoscape* [25]. Data taken from Tables S3 and S8 for full and reduced models, respectively.

7. Whitaker et al. [26]: 20 737 genes, enrichment performed on partial least squares component 2 using GORILLA [27]. Data taken from supplementary data table: `WhitakerVertes_PLSEnrichmentGeneList.xls`.

8. Romme et al. [28]: 20 737 genes, enrichment on top 100 genes with strongest correlation to SCZ connectome disconnectivity using ORA (*Panther* [19]). Data taken from supplementary tables.

9. Liu et al. [29]: 20 738 genes, biological process enrichment performed for: (i) left hippocampus (HCP atlas, data taken from Table S4), (ii) middle insular area (HCP atlas) (Table S5), (iii) brain-wide functional connectivity association study of autism (Table S7), and (iv) chronic schizophrenia (Table S8).

10. Kuncheva et al. [30]: 16 906 pre-selected genes, enrichment on clusters of spatial expression networks (SENs) for GO biological process (uncorrected threshold, $p < 0.001$) with categories reduced using *REVIGO*. Full results not provided; summaries taken from text.

11. Ritchie et al. [31]: 13 384 genes, enrichment on spatial correlation (Spearman's $\rho$) between expression patterns and the T1w:T2w ratio, using the AUROC method, ranked genes by correlation, including GO categories with between 10 and 200 genes after intersection with AHBA data (6885 GO categories). Data taken as $p_{\mathrm{FWER}}$ for biological processes: negative correlations (Table 2) and positive correlations (Table 3).

12. Diez and Sepulcre [32]: 3719 neural genes (determined from browsing *AmiGO*), enrichment on correlation between stepwise functional connectivity maps and cortical expression profiles, focusing *a priori* on brain-related categories. Enrichment performed as over-representation analysis on biological processes using *PANTHER* 13.1 at a threshold FDR $< 0.005$. Data taken from Supplementary Tables 3–6 as FDR-corrected $p$-values.

13. Wen et al. [33] analyzed for gene modules that are correlated to R2t$^*$ using *ToppGene*, with additional testing using *DAVID*. We noted GO Biological Process categories with $p < 0.05$ from the top 30 Annotation Clusters (up to an enrichment score of 1.27) from Dataset S3.

14. Liu et al. [34] analyzed how dynamic brain networks (analyzed from the viewpoint of the chronnectome) are spatially configured, and how these spatial maps are associated with gene transcription. We took data from Table S5 (Biological Process: PLS1), and Table S9 (Biological Process, PLS3).

15. Anderson et al. [35] analyzed enrichment for limbic network-biased genes using *ToppGene*. We took Biological Process results from the Supplementary Data 1 (sheet: `ToppGene_limbic_n505`).

16. Anderson et al. [36] analyzed enrichment for genes highly correlated to interneuron markers PVALB and SST using *ToppGene*. We took GO:BP data from Table 1.

17. Betzel et al. [37] used *GOrilla* to understand gene subsets that exhibit correlated expression patterns that are most strongly related to ECoG FC, providing results in frequency bands $1-4$ Hz (biological process results taken from Table S5) and $4-8$ Hz (biological process results taken from Table S7).

18. Meijer et al. [38] used *GOrilla* to find genes that are differentially expressed genes in the stress network (brain areas activated by stress in individuals with low or high stress sensitivity). Data was taken from Table S4.

19. Romero-Garcia et al. [39] performed PLS to find genes associated with changes in cortical thickness in autism, using *Enrichr* for enrichment in GO biological processes. Data was taken from the main text.

20. Liu et al. [40] investigated how emotion regulation and memory control related fMRI task activation maps correlate with gene expression. 1061 'inhibition-related genes' are correlated with all four tasks (memory control, emotion regulation, stop-signal, and go/no-go). Enrichment done using *GOATOOLS*. Data taken from Table S3.

21. Grothe et al. [41] analyzed differentially-expressed gene sets in Alzheimer's disease vulnerable brain regions using GSEA (gene set enrichment analysis) using all 20 737 genes (1036 GO-based gene sets were included). Enrichment results for GO biological processes were taken from Table 3 (differentially expressed gene sets in neurodegeneration-vulnerable brain regions).

22. Vidal-Pineiro et al. [42] investigated transcriptional patterns related to cortical thinning across the lifespan. GO-term enrichment was performed with VisuaL Annotation Display (VLAD). Data was taken from Table S1.

23. Yao et al. [43] proposed a method to perform enrichment by jointly considering gene sets (GS) and brain circuits (BC) to examine if a GS–BC pair is enriched in a list of gene–neuroimaging quantitative traits (QT, such as the average amyloid deposition). Enrichment results for GO biological processes were taken from Table 3.

### S4 Supplementary Figures

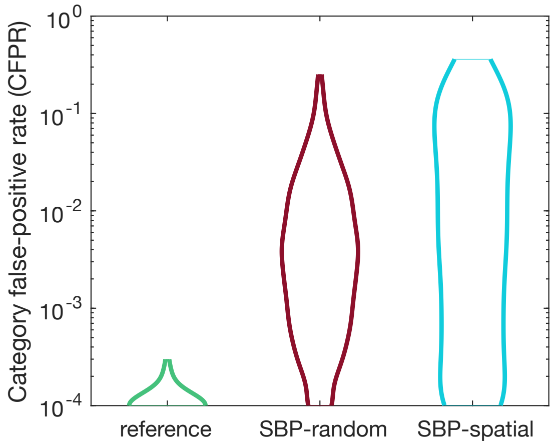


Figure S1: **Category-level false-positive rate (CFPR) across three null ensembles of 10 000 human cortical maps.** See Fig. 2A for information.

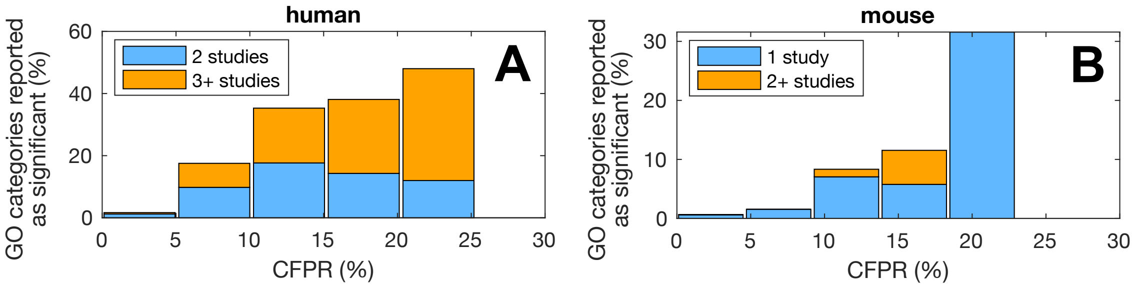


Figure S2: Across five equally-spaced bins of CFPR, bars show the percentage of GO categories that have been reported as significant in the literature survey of **A** human, and **B** mouse. The larger number of human studies allowed us to distinguish between GO categories reported in two, or three or more studies; in mouse we distinguish between one, or two or more studies.

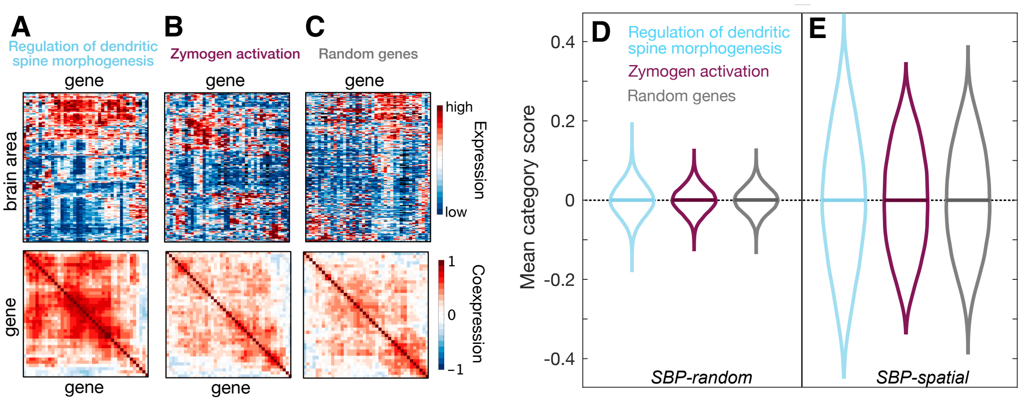


Figure S3: **GO categories with high within-category gene–gene coexpression are most prone to false-positive enrichment**. We plot expression (brain area × gene) and coexpression (gene × gene) heat maps for two example GO categories in the mouse brain: **A** 'regulation of dendritic spine morphogenesis' (40 genes); and **B** 'zymogen activation' (42 genes); as well as **C** a random set of 40 genes, for comparison. Each gene's expression is normalized (low to high) for visualization purposes. Genes annotated to 'regulation of dendritic spine morphogenesis' display a more characteristic spatial patterning and hence have higher coexpression. Distributions of each category's score (mean correlation between the genes in that category and a phenotype) across an ensemble of null phenotypes, are plotted as violin plots for: **D** the SBP-random ensemble of random-number phenotypes (cf. Fig. 2A(ii)), and **E** the SBP-spatial ensemble of random spatially autocorrelated phenotypes (cf. Fig. 2A(iii)). The mean of each distribution is annotated with a horizontal line.

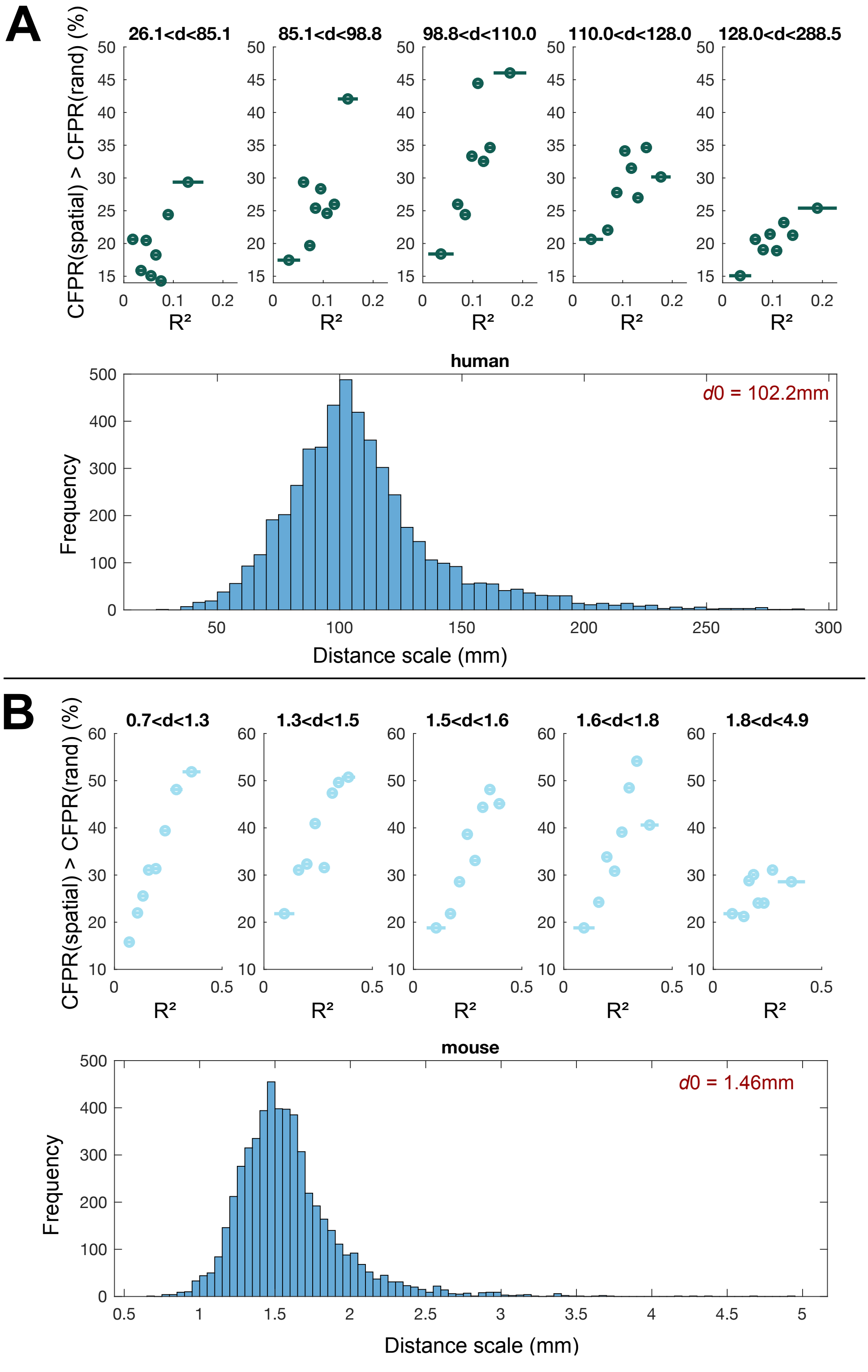


Figure S4: **CFPR (SBP-spatial) > CFPR (SBP-rand) is greatest for GO categories with strong spatial autocorrelation ($R^2_{\text{exp}}$) and a similar distance scale of autocorrelation.** For **A** human, and **B** mouse, we show a distribution of fitted distance scales, $\lambda$, across all GO categories. For each category, this was estimated from an exponential fit to correlated gene expression (of a given category of genes) as a function of distance. The global value, $d_0$ (obtained from including all genes in the correlated gene expression calculation, and used to construct the SBP-spatial ensemble) is annotated in the upper right of the distribution plot. Across equiprobable bins of $R^2_{\text{exp}}$ and for a given distance range, upper plots show the percentage of GO categories in each bin that display an increase in CFPR under the SBP-spatial null ensemble relative to the SBP-random null ensemble (as Fig. 3B).

### S5 Supplementary Tables

| GO Category | Mouse CFPR [SBP-spatial] (%) | Human CFPR [SBP-spatial] (%) | References |
|---|---|---|---|
| Regulation of synaptic plasticity | 23 [37] | 10 [7] | [11, 20, 29, 34, 38] |
| Regulation of postsynaptic membrane potential | 19 [35] | 10 [17] | [11, 20, 32] |
| Glutamate receptor signaling pathway | 16 [34] | 6 [14] | [12, 20, 29, 43] |
| Respiratory electron transport chain | 12 [4] | 20 [30] | [11, 20–22, 26] |
| Learning | 20 [34] | 5 [2] | [16, 20, 29, 38] |

Table S1: **GO categories with high category false-positive rates (CFPRs) are predominantly related to neuronal and metabolic biological function in mouse and human.** We list selected GO categories with amongst the highest CFPRs across mouse and human, across ensembles of spatially independent random phenotypes (SBP-random) and spatially autocorrelated random phenotypes (SBP-spatial). CFPR is listed for SBP-random [and for SBP-spatial in brackets]. A full ordered list is in Supplementary File `CFPRTable.csv`.
